# Accounting for genetic interactions improves modeling of individual quantitative trait phenotypes in yeast

**DOI:** 10.1101/059485

**Authors:** Simon K. G. Forsberg, Joshua S. Bloom, Meru J. Sadhu, Leonid Kruglyak, Örjan Carlborg

**Author notes:** Correspondence should be addressed to Ö.C.

## Abstract

Experiments in model organisms report abundant genetic interactions underlying biologically important traits, whereas quantitative genetics theory predicts, and data support, that most genetic variance in populations is additive. Here we describe networks of capacitating genetic interactions that contribute to quantitative trait variation in a large yeast intercross population. The additive variance explained by individual loci in a network is highly dependent on the allele frequencies of the interacting loci. Modeling of phenotypes for multi-locus genotype classes in the epistatic networks is often improved by accounting for the interactions. We discuss the implications of these results for attempts to dissect genetic architectures and to predict individual phenotypes and long-term responses to selection.

---

When the combined phenotypic effect of alleles at two or more loci deviates from the sum of their individual effects, it is referred to as a genetic interaction or epistasis. Most biological traits are regulated by a complex interplay between multiple genes and environmental factors. Despite this underlying complexity, data and theory have shown that it is expected that most of the genetic variance in a population will be additive^1–3^. The apparent contradiction between the complexity of the biological mechanisms that determine quantitative traits, and the observation that most genetic variance can be captured by an additive model, has led to a long-standing debate in genetics: Does the predominant role of the additive genetic variance mean that strictly additive models are always sufficient to describe the relationship between the genotype and the phenotype of an organism^1,4^, or could there be added value in explicitly modeling genetic interactions despite the lower levels of epistatic genetic variance^5–8^?

There are some situations where data and theory has suggested that it might be particularly important to account for genetic interactions. One is when the aim is to predict phenotypes of individuals based on their genotype. If interactions lead to extreme phenotypes for some genotypes, these phenotypes are unlikely to be captured by additive models, particularly if they are rare. This has, for example, been illustrated for sporulation efficiency in yeast^9^. Another is in the prediction of long-term selection response. Under additivity, both the additive variance and the response are expected to be near constant over the first few generations. As generations proceed, allele frequencies change to alter the additive variance and consequently the response to selection. This change is more rapid for traits regulated by fewer loci with larger effects than for traits regulated by many loci with smaller effects. It is known that genetic interactions can contribute to the additive genetic variance in a population^1,7^. The contribution, however, varies depending on the joint allele frequencies across all the interacting loci as well as on the types and strengths of the genetic interactions^10,11^. The changes in the additive variance, and hence the response, during ongoing selection is therefore more complex in the presence of genetic interactions. As a result, genetic interactions can make the long-term selection response more dynamic^12,13^ and result in a realized response beyond predictions based on the additive genetic effects and allele frequencies at the individual loci^10,11^. However, as little is known about how prevalent and strong genetic interactions are in real populations, and how much they contribute to the additive variance as the allele frequencies change during selection, it has been difficult to obtain any empirically based conclusions about how influential interactions are expected to be in these situations.

Here we analyze a panel of 4,390 yeast recombinant offspring (segregants) from a cross between a lab strain (BY) and a vineyard strain (RM), generated in Bloom, *et al.* 2015^3^. In this population, each segregant is genotyped for 28,220 SNPs and phenotyped for 20 end-point growth traits. Across these traits, a total of 939 QTL with additive, and 330 with epistatic, effects were mapped previously^3^. Since the individuals in this population are haploids, the sample size is large and all allele frequencies are close to 50%, this dataset presents a unique opportunity to accurately estimate how allelic combinations across large numbers of loci influence quantitative traits. Using phenotype information for multiple segregants with each possible allelic combination across many loci, we directly estimate how high-order genetic interactions contribute to complex trait variation in this segregating population. We quantify how well quantitative genetics models can capture the empirically revealed relationships between multi-locus genotypes and phenotypes.

We observe networks of epistatic loci for most of the traits in the study, and find that some highly interactive loci in these networks can hide, or reveal, the effects of their interactors. We show that additive genetic models capture much of the genetic variance contributed by the interacting genes in these networks. However, when used to estimate the phenotypes for individual segregants, they often fail to fully capture the effects of multi-locus genotypes that lead to extreme phenotypes. Accounting for interactions in the models led to more accurate phenotypic predictions for such genotypes. We illustrate this by analyzing an individual network in detail, and then generalize the results across the other revealed networks. We discuss the potential impact of these findings on prediction of individual phenotypes that is of importance in for example personalized medicine, prediction of long-term selection response in breeding and evolution, and interpretation of results from QTL and genome-wide association studies.

## Results

### Many epistatic QTL are part of highly connected networks

Most epistatic QTL^3^ interacted with one or a few loci, while a smaller subset were involved in pairwise interactions with several loci (Figure 1A). By visualizing the 330 statistically significant epistatic QTL as nodes, and the interactions between them as edges, we revealed many networks of interacting loci. These were often connected in hub-and-spoke type of architectures where QTL involved in many interactions tied larger networks together (Figure 1B). We refer to these as radial networks, with a hub-QTL in the center that connects the radial QTLs. Hub-radial QTL interactions were, on average, more significant than interactions that did not involve a hub (Supplementary Figure 1), supporting that the radial architecture is a prominent feature of the networks. The available genotype and phenotype data allowed us to accurately estimate the phenotypes for individual six-locus genotype-classes (see below). We therefore selected the 15 six-locus radial networks where a hub-QTL interacted with at least five other QTL, for further in-depth studies (Figure 1B). The selected networks contributed to 11 of the 20 studied traits, and included 81 QTL.

**Figure 1.**
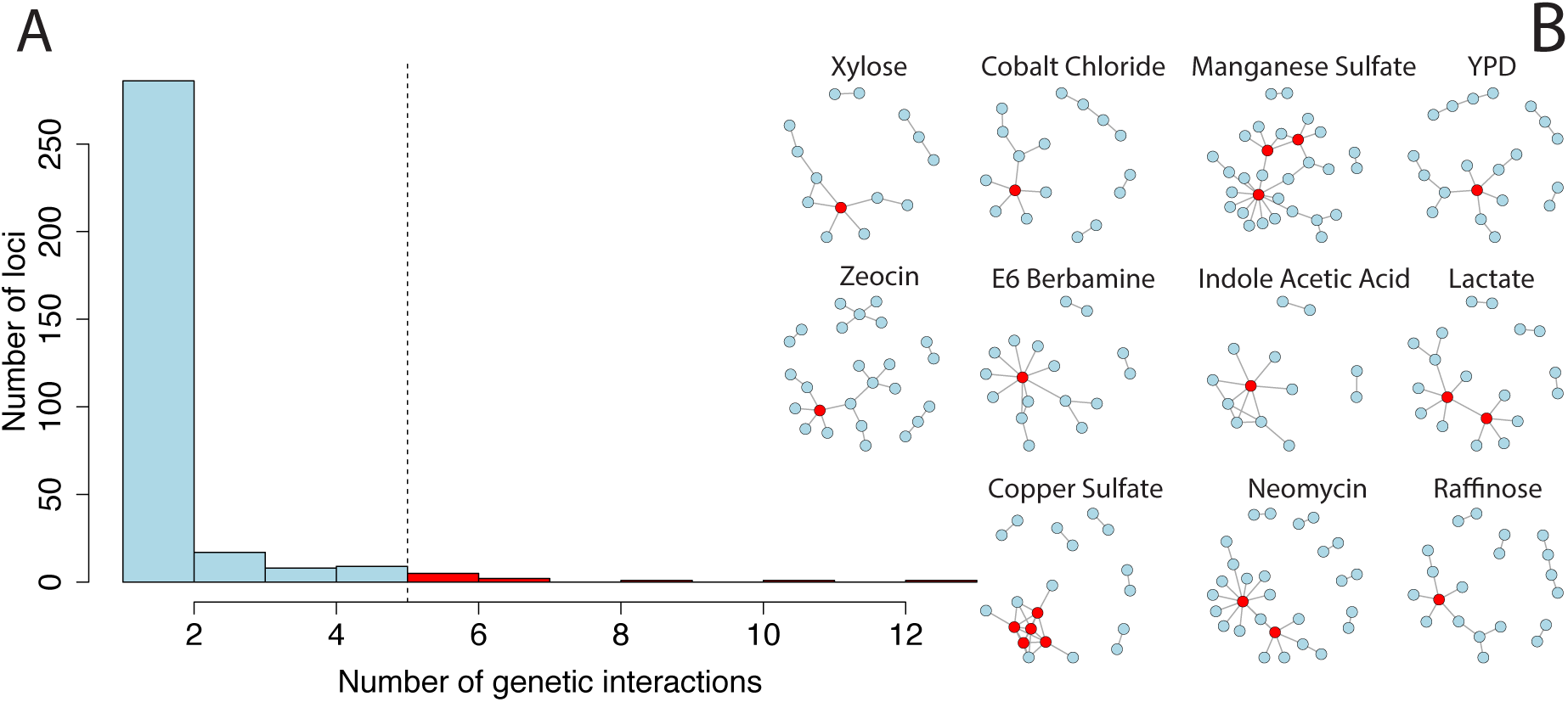
Many QTL involved in pairwise interactions are part of highly interconnected epistatic networks. **A)** A histogram of the number of interactions each epistatic QTL is involved in. Most of the 330 epistatic loci detected for the 20 traits are involved in few pairwise interactions. Few QTL are involved in many interactions, here defined as five or more, but their role is prominent since they are the hubs that tie the networks together **(B). B)** The pairwise QTL interactions contributing to growth in 11 different media form highly interconnected epistatic networks. Each circle represents a QTL and the lines represent significant pairwise interactions^3^. Red circles highlight hub-QTL involved in five or more pairwise interactions.

Below, we first illustrate our analyses and results for the network regulating growth on medium containing indoleacetic acid (IAA-network) and then extend them to all networks.

### Hub-QTL alleles often moderate the phenotypic effects of radial QTL alleles

The hub-QTL in the IAA-network explained the most additive genetic variance of any locus for this trait 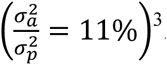. The phenotypic variance was 3-fold higher among segregants with the BY-genotype, than among segregants with the RM-genotype, at this locus. This genetic variance-heterogeneity is highly significant (*p* < 1×10^−16^). By estimating the narrow sense heritability (h^2^) separately among segregants with the *BY* and *RM* allele at this hub-QTL we showed that much of this difference was due to genetics 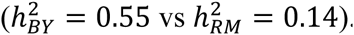. We here call such QTL, where one allele suppresses genetic contributions by other loci and the other allele uncovers them, genetic capacitors. Across the 15 epistatic networks, 10 hub-QTL were genetic capacitors with significant differences in h^2^ between the genotypes ranging from 10 *to* 42%. By testing 40 randomly selected radial QTL, we found that few (3) of these were significant genetic capacitors and that they were weaker capacitors than the hubs (mean difference±SD in *h*^2^for radial QTL = 4.3%±3.6 vs 12.5%±9.2% for hub-QTL). We also revealed a strong correlation (*r*^2^ = 0.64; *p* < 1×10^−16^) between the level of variance-heterogeneity between the genotypes at the 330 epistatic QTL and the number of interactions they were involved in (Supplementary Figure 2). Together, this shows that strong genetic capacitors^14,15^ often are hubs in epistatic networks.

**Figure 2.**
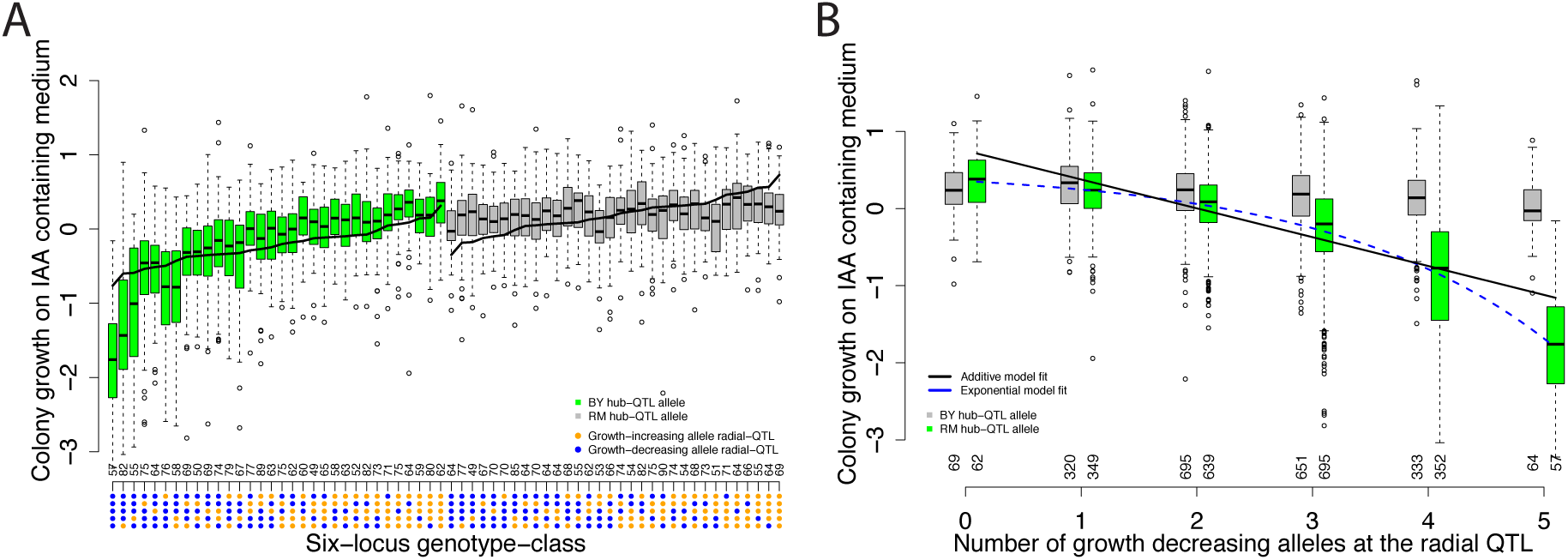
An epistatic network regulates growth on indoleacetic acid containing medium. One allele at the network hub-QTL hides the phenotypic effects of segregating alleles at the radial QTL. When the other allele is present, the growth decreasing effects of the radial QTL are combined in a non-linear fashion. Tukey boxplots illustrating the phenotypic distributions in segregants with different combinations of alleles across the IAA-network. **A)** One boxplot for each of the 64 genotype classes, where the color indicates the genotype at the hub-QTL (chrVIII:114,114bp; Green/grey boxes for BY/ RM alleles). The x-axis gives the six-locus genotype class, where blue/orange dots indicates growth-decreasing/increasing alleles at the five radial-QTL (from top to bottom chrXIV:469,224bp, chrIII:198,615bp, chrIV:998,628bp, chrXIII:410,320bp, and chrXII:645,539bp). Genotypes with an RM allele at the hub-QTL have less variable (canalized) phenotypes, whereas genotypes with a BY allele have more variable (capacitated) phenotypes. The black line through the boxes illustrates the additive model based estimates of the phenotypes for the 64 genotype classes. The number above the x-axis is the number of segregants in each genotype class. **B)** One boxplot for each group of segregants that share the same number of growth decreasing alleles at the five radial QTL. The segregants are divided and colored based on the genotype at the hub-QTL as in A). The x-axis gives the number of growth decreasing alleles at the radial QTL and the number of segregants in each group. The regression lines illustrate the fit for linear additive (black; R2 = 0.34) and non-linear exponential (dashed blue; R2 = 0.39) models, respectively.

### Deviations between estimated and modeled phenotypes for individual multi-locus genotype-classes

#### The contribution by allelic interactions to complex trait variation in a segregating population

For the six-locus IAA-network, we divided the segregants into 64 groups representing each of the 64 possible six-locus genotype classes, and calculated their phenotypic means and variances (Figure 2A). Segregants with the capacitating *BY* allele at the hub-QTL (green) display the poorest growth when they have many IAA-sensitizing alleles at the five radial QTL. In contrast, segregants with the canalizing *RM* allele at the hub-QTL (grey) have similar growth regardless of how many IAA-sensitizing alleles they have at the radial QTL. The *RM*/*BY*alleles at the hub-QTL in the network thus decrease (canalize)/increase (capacitate) the effects of the radial QTL, respectively (Figure 2A). Similar results are also observed for several of the other networks with significant differences in h^2^ between the genotypes at the hub-QTL (Supplementary Figure 3).

Across the networks, we detected hub-QTL capacitor alleles of both *BY* and *RM* origin. The most extreme phenotypes in these networks were always observed for a genotype-class with a combination of *BY* and *RM* alleles at the hub and radial loci. The alleles at the radial loci that required the presence of the capacitating hub allele to reveal their full effect on growth (Figure 3; Supplementary Figure 3; Supplementary Figure 4) did in 60% of the cases originate from the same strain as the canalizing hub allele. The two parental strains thus harbor cryptic, or hidden, genetic variation^16–19^, whose phenotypic effect is revealed when combined in the haploid segregants.

**Figure 3.**
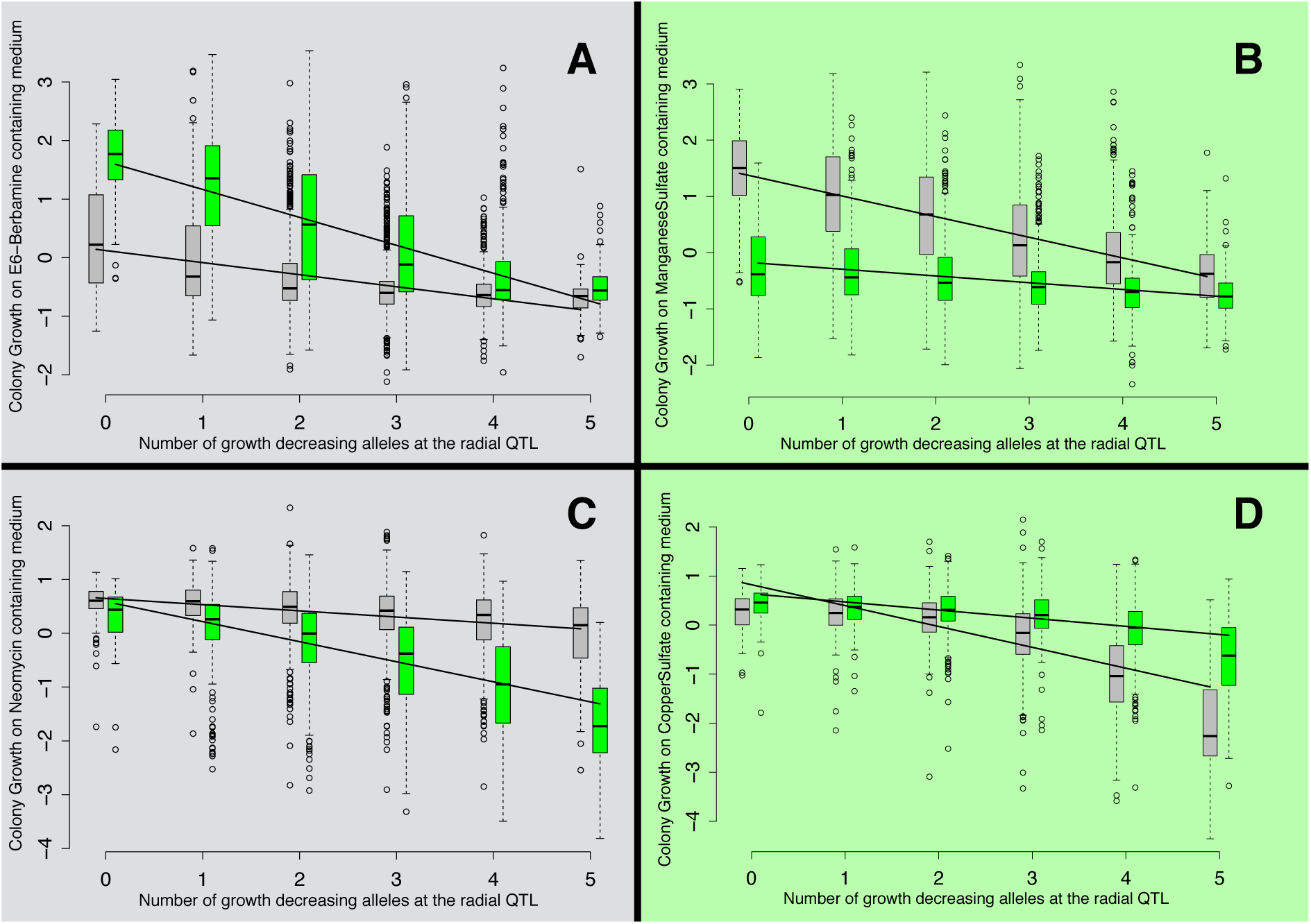
The epistatic networks contain hub-QTL capacitor alleles of both BY and RM origin. In some networks they moderate growth increasing, and in others growth decreasing, effects of segregating alleles at radial QTL. The boxplots show phenotypic distributions for groups of segregants with different numbers of growth affecting alleles at the five radial QTL in six-locus epistatic networks. Each sub-figure corresponds to one epistatic network where the hub-QTL capacitates the growth-increasing, or growth-decreasing, effects of segregating alleles at the radial QTL (**A**: E6-Berbamine, **B**: Manganese Sulfate, **C**: Neomycin, **D**: Copper Sulfate). Within each network, the segregants are divided based on the genotype at the hub-QTL, and segregants carrying the RM allele are shown in grey and those with the BY allele are shown in green. The x-axis gives the number of growth decreasing alleles at the radial QTL. **A(B)** illustrate networks where the RM(BY) allele at the hub-QTL maintain low growth in the segregants almost regardless of the genotypes at the radial-QTL. In **C(D)**, the RM(BY) maintain high growth almost regardless of which alleles segregate at the radial QTL. The regression lines illustrate the difference in effects of the radial-QTL alleles depending on the genotype at the hub-QTL.

#### Non-linear capacitation effects in some epistatic networks

In the IAA-network, the reduction in growth among the segregants with the *BY* allele at the hub-QTL decelerates in a multiplicative, rather than a linear, manner as the number of IAA-sensitizing alleles increases (Figure 2B). For segregants with the *BY* allele at the hub-QTL, the effect of having 5 IAA-sensitizing alleles is much larger than 5 times the effect of having one IAA-sensitizing allele. As a result, an exponential model fits this data better than an additive model (R2 increases from 0.34 to 0.39; Figure 2B). One other network, affecting growth in medium containing Copper Sulfate, displayed a similar non-linear capacitation (R2 increased from 34 to 43%; Figure 3D; Supplementary Figure 4). This multiplicative effect could result from measuring growth as the increase in radius of the yeast colonies, or it could be a feature of the underlying biology.

#### Extreme additive-model based estimates of individual phenotypes are often biased

Many additive-model based estimates of the phenotypes for multi-locus genotype classes in the IAA network differed substantially from the actual values estimated directly from the data (Figure 2A). Cross-validated model-based estimates were computed for each multi-locus genotype class to quantify their accuracy and bias. Accuracy for each genotype class was measured by the mean square error (MSE), and bias by the difference between the modeled and actual phenotypes. Twenty-three of the 64 estimates were significantly biased (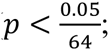 two-sided t-test), showing that the additive model was unable to represent the genetic contributions by the IAA network to many of the individual segregant phenotypes. To evaluate whether this trend generalized across all networks, and whether alternative quantitative genetics model parameterizations could perform better, we fitted three different quantitative genetics models to all six-locus networks: i) additive effects only, ii) additive effects and pairwise interactions and iii) additive, pairwise and three-way interactions.

Models with only additive effects captured much of the phenotypic variance for all networks (on average 28% of 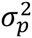). Accounting for epistatic interactions only increased the variance explained marginally (on average 5% of 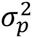). The additive model based estimates of the phenotypes were, however, significantly biased for between one and 23 of the 64 genotype classes per network in 14 of the 15 examined networks (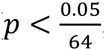; two-sided t-test; Figure 2; Figure 4; Supplementary Figure 5). Models with pairwise interaction terms provided unbiased estimates of all 64 measured genotype values for most networks. Only two networks required three-way interaction terms to remove all detectable bias. In 5 of the 15 networks, the accuracy was significantly better for at least one of the 64 genotype classes when using models with pairwise interaction terms (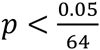; two-sided t-test; Supplementary Figure 6).

**Figure 4.**
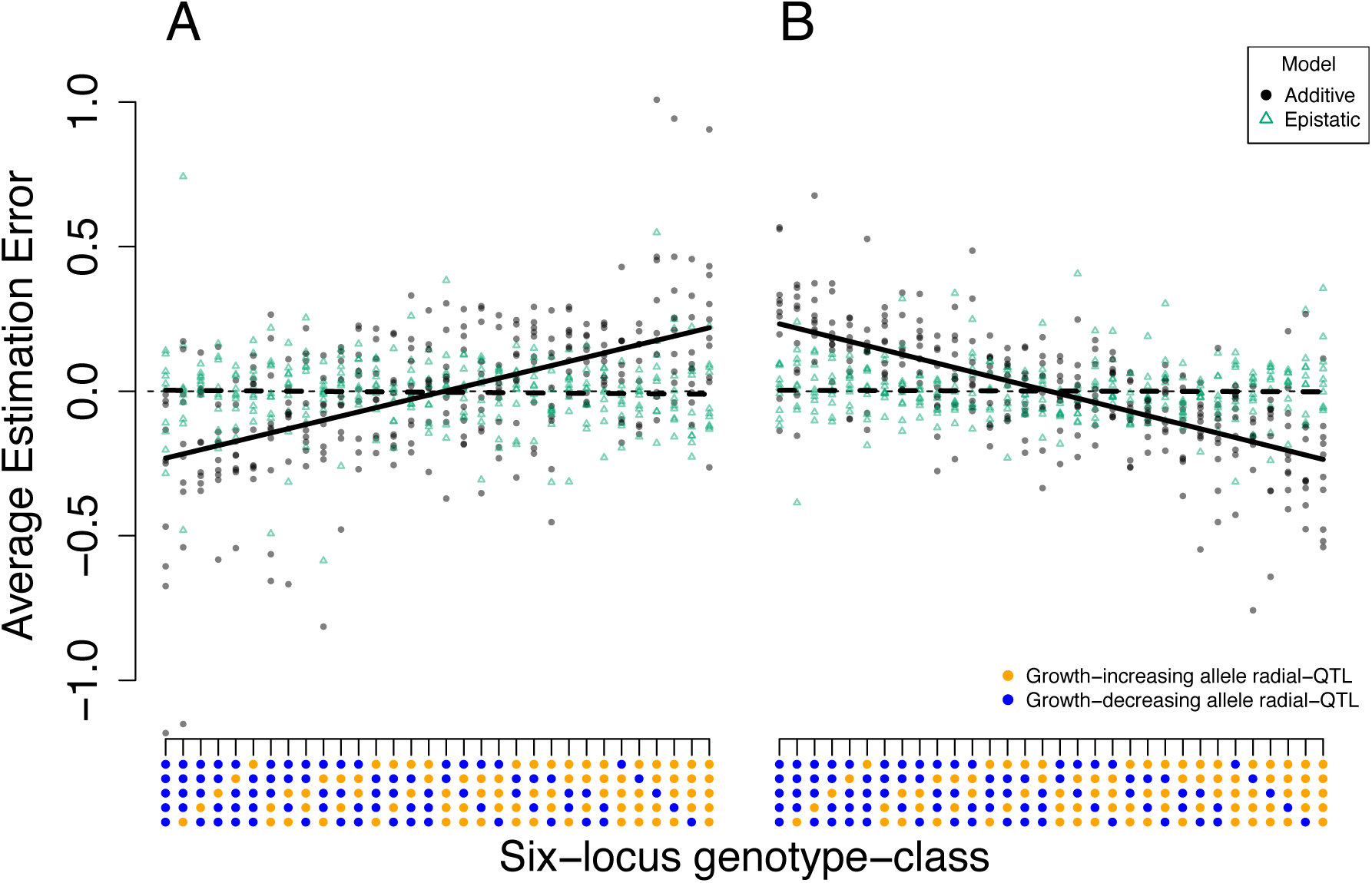
The biases in the additive model based estimates of the phenotypes are largest in the genotype-classes with the largest or poorest expected growth. The y-axis gives the average estimation-error (bias) from cross-validation for each genotype in the 10 networks. The x-axis illustrates the genotype at the five radial QTL in the networks with blue/orange dots indicating growth-increasing/decreasing alleles respectively. Each dot represents the average estimation error (bias) for a particular six-locus genotype-class in one of the 10 networks. Black dots correspond to models with only additive effects and green triangles to models with both additive effects and pairwise interactions. The estimation errors (biases) are calculated as (measured – estimated) phenotypes, and over/under estimations will therefore result in negative/positive estimation errors (biases) respectively. Across the 10 interaction networks with a capacitor hub-QTL, the most extreme additive model based representations of individual multi-locus genotypes are biased. There is a positive correlation between the estimation errors (bias) and the number of growth-increasing alleles at the radial-QTL when there is a capacitor allele at the hub-QTL (**A**; r = 0.54; p < 10^−8^). The additive model thus underestimates the extremity of the genotypes with many growth-decreasing alleles at the radial-QTL, and overestimates the extremity of the genotypes with many growth-increasing alleles, at the radial-QTL. An opposite trend (r = -0.69; p < 10^−8^) is observed among segregants with a non-capacitor allele at the hub-QTL (**B**). There, the additive model over- (under-) estimates the extremity of the segregants with many growth-decreasing (increasing) alleles at the radial-QTL

The bias for the additive model estimates of the phenotypes of individual multi-locus genotypes was largest for the most extreme trait values, i.e. those corresponding to the best or worst growth in the capacitated group, and the best or worst estimated growth in the canalized group (Figure 2; Figure 4). In networks where the hub-QTL are capacitors, the direction of the bias depends on the genotype at the capacitor: Among segregants with the capacitor allele at the hub-QTL, the models underestimated the phenotypic effect of combining many growth-increasing or growth-decreasing alleles at the radial-QTL (Figure 4A). For segregants with the canalizing allele at the hub-QTL, the models instead overestimated these phenotypic effects (Figure 4B). By accounting for epistatic interactions, the bias is reduced or entirely removed (Figure 4). For the 5 networks where the hub-QTL was not a significant capacitor, the bias was not dependent on the genotype at the hub-QTL.

### The genetic variances explained by the epistatic networks are highly dependent on allele frequencies

The additive genetic variance contributed by a locus depends on its effect and allele frequency in the analyzed population^20^. When loci interact, the variance explained marginally by each of the epistatic loci will in addition also depend on the frequencies at the loci with which it interacts. For example, the additive genetic variance contributed by each individual locus in the IAA-network varies depending on the allele frequencies of all loci in the network due to the extensive interactions among them. When considering the entire population of segregants (all allele frequencies ~0.5), the additive variance (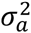) contributed by the network amounts to 26.8% of the total phenotypic variance in the population. In the subpopulations where the *BY*/*RM*alleles at the hub-QTL are fixed (*BY* allele frequency 1 or 0), the network instead contributes 36% and 3% of the total phenotypic variance, respectively. To generalize this result across the allele-frequency space, we simulated populations with allele frequencies ranging from 0.05 to 0.95 at increments of 0.15 for the six loci in the network. We then evaluated how the additive genetic variance contributed by the individual loci varied depending on the allele frequencies at the other five loci. We also simulated populations without genetic interactions. In Figure 5 we summarize the results for the simulations based on the 64 actual genotype-values in the IAA-network. The additive genetic variance contributed by the hub-QTL varied from 0 to 58% of the total phenotypic variance in the population, only by changing the allele-frequencies at the five other loci (Figure 5). The result was similar for the other five QTL, although their ranges were smaller than for the hub-QTL (Figure 5 blue boxes; average range: 0 to 31% of 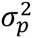). As expected, the additive genetic variance contributed by each locus was much less dependent on the allele frequencies when loci with the same marginal effects, but no interactions, were simulated (Figure 5 red boxes; range: 2 to 6% of 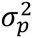). The results were similar across all 15 networks, with estimates of additive genetic variance for individual epistatic loci that were highly dependent on the allele frequencies at the other loci in the network.

**Figure 5.**
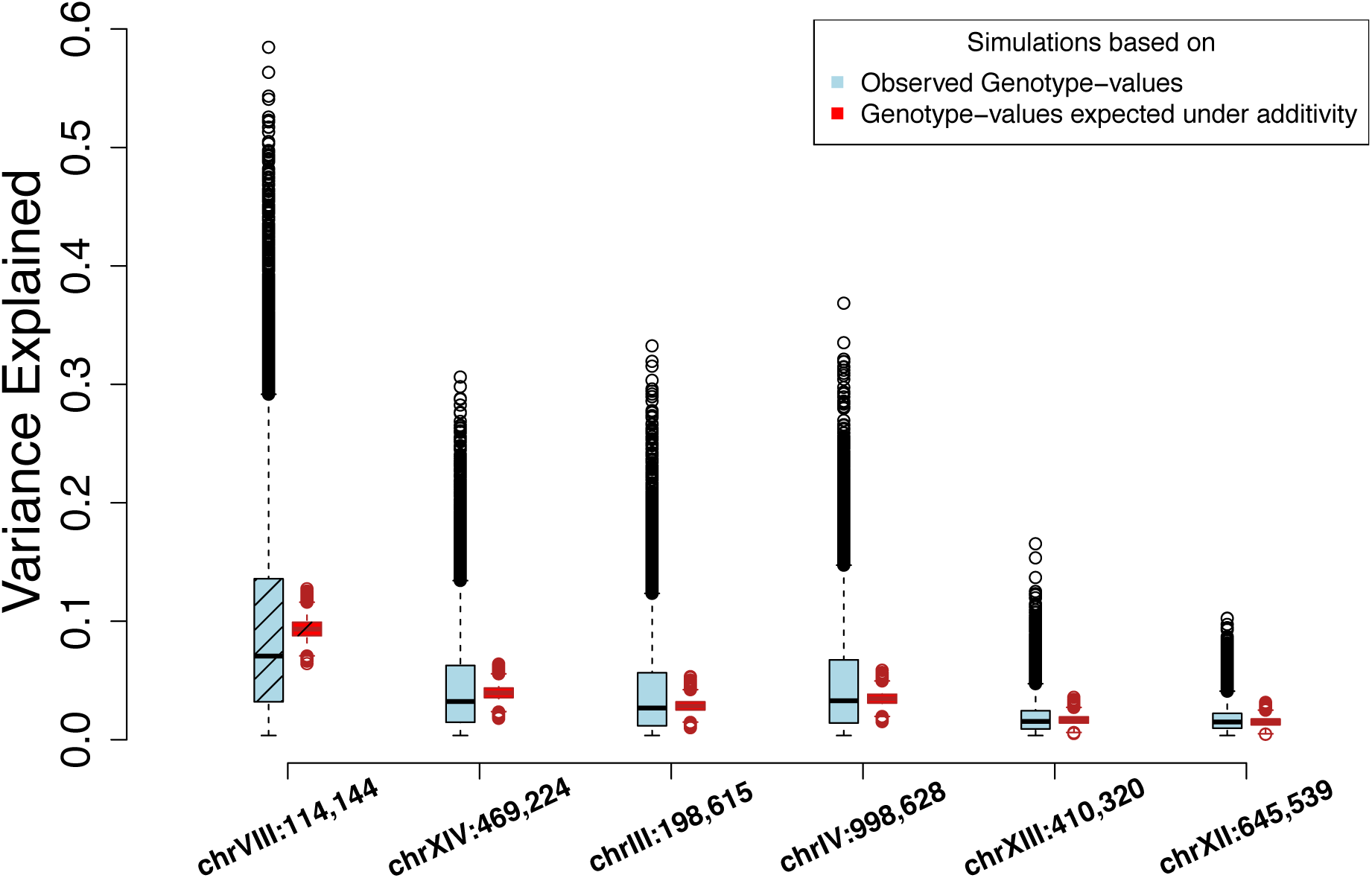
Simulations show that the additive genetic variances contributed by the loci in the epistatic network regulating growth on indoleacetic acid containing medium is highly dependent on the allele-frequencies at the other loci in the same network. The phenotypic variance explained by the individual loci in the IAA-network depends on the allele frequencies at the other loci with which they interact. Each boxplot shows the additive genetic variances across 16,807 simulated populations. For each locus, we simulated populations where the allele frequency of the loci themselves were fixed at 0.5, while varying the frequencies at the other 5 loci from 0.05 to 0.95 in increments of 0.15. The light blue boxes represent the variability in the additive genetic variances when simulating populations based on the observed phenotypic means for the 64 genotype classes in the IAA-network. The red boxes represent the additive genetic variances when simulating based on the phenotypic means estimated using the additive genetic model (black line in Figure2A). The x-axis gives the location of the SNP representing each locus (chromosome:location in bp). The two leftmost boxplots (diagonal lines) show the simulation results for the hub-locus.

## Discussion

The link between the genotype and the phenotype of an organism is immensely complex. Despite this it can, to a great extent, be captured using models that assume that gene variants combine their effects in an additive manner. We here used a large experimental yeast cross to identify six-locus epistatic networks affecting 11 complex traits. We then estimated the average phenotypes for the groups of segregants sharing each combination of alleles at these loci. We evaluated how well different quantitative genetics models captured the phenotypes of these multi-locus genotype classes. In most networks, the phenotypes for at least one, but often several, multi-locus genotype classes deviated significantly from what is expected under additivity. This empirically illustrates the important role of classic epistasis, as defined by Bateson more than 100 years ago^21^, in the genetic architecture of complex traits. We provide several examples of such epistasis, involving multiple loci in highly interconnected genetic networks.

An earlier study of this population^3^ showed that most of the genetic variance for the 20 measured traits is additive. Consistent with this, the additive model based estimates of the phenotypes for most multi-locus genotype classes in the epistatic networks were reasonable. However, the most extreme estimates from the additive models were often both inaccurate and biased (Figure 4). For example, the bias is very large for the most extreme genotype class in the IAA-network (1.7 *σ*_*p*_; Figure 2). Our results highlight the importance of analyzing collected data with models that can represent the features of the underlying genetic architectures. They also confirm that, regardless of the underlying genetic architecture, additive models are likely to capture much of the genetic variation for a trait. This makes them useful for revealing genes contributing to the phenotypic variance in a particular population, as well as for predicting short-term response to selection in a population^11^. However, we also show that additive models are often unable to represent all key features of genetic architectures involving networks of epistatic loci. In particular, their most extreme estimates are often inaccurate and biased for networks with capacitor hub-QTL. Accounting for epistasis increases estimation accuracy and decreases bias. Modeling genetic interactions should therefore be considered when it is important to identify and predict the effects of specific combinations of alleles, or where it is important to identify genotypes that are likely to lead to extreme phenotypes. Examples of this include prediction of disease risk or drug responses in individual patients.

As shown here and in earlier studies^1,7,22^, most genetic variance in a population is expected to be additive even in the presence of extensive epistasis. The lack of empirical knowledge about the pervasiveness and strength of epistasis in the genetic architectures of complex traits makes it largely unknown how much of the observed additive genetic variance in quantitative genetics studies is due to genetic interactions. This experimental yeast population allowed us to directly estimate the phenotypes for individual multi-locus genotype classes in networks of interacting loci. With these as a basis, we used simulations to demonstrate that the types of interactions revealed here have a very large influence on both the estimates of the additive genetic effects of the individual loci and their contributions to the additive genetic variance. The IAA-network provides a striking example: the additive variance contributed by the hub in the epistatic network (Figure 2) ranged from zero to the largest contribution by any single locus across all networks when we varied the allele frequencies at the 5 radial QTL. This empirically illustrates how allelic interactions (epistasis) can be the main driver of the additive genetic variance in a population, and that the importance of epistasis in the genetic architecture of a complex trait cannot be inferred from the relative levels of additive and epistatic genetic variance.

Many interacting loci in this population were part of radial epistatic networks where hub loci interact with multiple other QTL. In general, this network topology reflected how the loci contributed to the phenotypic variation in the population. The hub-QTLs acted as genetic capacitors that modify the effects of the radial loci in the network. These capacitating interactions are highly influential for the total level of phenotypic variance displayed in a population, as they can both buffer and release cryptic (standing) genetic variation^23,24^. Several genetic capacitors have been studied in molecular detail, including the heat shock protein *HSP90*^16,25^ and *EGFR*^26^. Genetic capacitation has also been found to facilitate extreme selection responses, for example in a long-term experimental selection experiment in chicken^10–12^. The finding that genetic capacitor networks are common and influential for many traits in this population suggests that they should be considered in other studies of complex traits, including those aiming to genetically dissect, or statistically predict, responses to long-term selection.

It is currently unknown how indoleacetic acid affects yeast fitness, but the discovery of the epistatic network described here may shed some light on its mode of action. The hub-QTL in the IAA-network maps to the gene *GPA1,* which is required for the yeast response to mating pheromone. Although this response is not normally triggered under laboratory growth conditions, as the yeast is not exposed to mating pheromone, the *BY* allele of *GPA1* leads to residual expression of the pheromone response pathway^27,28^. Thus, a model for the capacitance activity of the *BY* allele of *GPA1* is that indoleacetic acid primarily affects cells with an activated pheromone response pathway. Interacting radial QTL would then arise if the underlying variants influence either the response to indoleacetic acid, or the activation of the pheromone response pathway by *GPA1*_*BY*_. The radial QTL include several genes involved in pheromone response, including *MAT*, which dictates which pheromones are expressed or sensed (and is known to interact with *GPA1*)^29^, and *VPS34,* which is required for *GPA1*’s activation of the mating pathway^30^. Further work will be required to elucidate the importance of the yeast pheromone response pathway for the fitness effects of indoleacetic acid.

For several other networks, the hub-QTL maps to candidate genes with known polymorphisms in BY and RM. For example, the networks regulating growth on media with Copper Sulfate and Manganese Sulfate have hub-QTL that maps to *CUP1* and *PMR1*. The *CUP1*_*BY*_-allele^31^ is known to increase copper ion tolerance^32^ and the *PMR1*_*RM*_ allele confers manganese resistence^33^. The current data does not allow further functional dissection of the possible connections between these polymorphisms and the capacitation in the networks. However, by revealing the allelic dependencies between the loci in the network (Figure 3B; Figure 3D), it is possible to formulate hypotheses about how these and other known polymorphisms in hub-QTL could contribute epistatically to growth in the respective media for testing in future functional studies (Supplementary text).

In summary, we show that networks of capacitating genetic interactions are common, and that these networks form a key part of the genetic architectures of multiple complex traits in a large experimental yeast population. We illustrate how such interactions affect model-based estimation of individual phenotypes and the inference of genetic architectures. This shows that epistasis needs to be explored beyond estimates of epistatic genetic variances, in order to understand its contribution to the phenotypic variability and long-term selection responses in populations. This is a key discovery in the long-standing debate about how to approach epistasis in complex trait research.

## Online Methods

The creation of the BYxRM cross, genotyping, phenotyping, quality control of genotypes, filtering and normalization of growth measurements has previously been described in^2,3^. The reanalyzed data are available as supplementary information in^8^. Additive QTLs were mapped in Bloom *et al.*^3^. All analyses were performed using the R framework for statistical computing^34^. All figures were prepared using R.

## Statistical analysis

### Inferring epistatic networks

Pairwise epistatic interactions were mapped by Bloom *et al.*^3^. Networks of epistatic loci were inferred by connecting loci that displayed pairwise interactions. The R-package *igraph*^35^ was used to visualize individual networks and to identify network hubs. The GWA analysis for growth on indoleacetic acid containing medium among the segregants with the *BY*-allele at the hub-locus was performed using the *qtscore* function in the R-package *GenABEL*^36^. Genome-wide significance was determined using a Bonferroni-corrected significance threshold for the number of tested markers 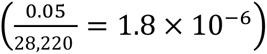. The additive genetic variance explained by a certain set of QTL was calculated as the R^2^ from a fixed effect model without interactions.

### Exhaustive mapping of loci in the network affecting growth on indoleacetic acid containing medium

To identify all individual loci that contributed the additional genetic variance in amongst the segregants carrying the *BY* allele at the hub-QTL in the IAA-network, we performed a linkage analysis in this group of segregants. This revealed in total 8 genome-wide significant loci in the radial network, out of which 6 were the same as the loci in the earlier two-way interaction analysis^3^.

### Estimating average phenotypes for multi-locus genotypes

The average phenotypes were estimated for each of the 2^6^ = 64 possible combinations of alleles for 15 six-locus epistatic networks. Each of these networks had a hub-QTL connected to five radial loci by pairwise interactions. On average, 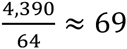 segregants are expected in each six-locus genotype class in these networks, allowing confident estimation of the average growth associated with carrying each possible combination of alleles at these loci. Some of the hub-QTL were connected to more than five other loci in the network in the initial network analysis. Here we only kept the loci with the strongest statistical interaction with the hub-QTL as we could not confidently estimate phenotypes for individual genotype-classes in networks with more than six loci.

### Estimation of the genetic variance heterogeneity at a locus

We estimated the difference in the phenotypic variance between segregants that carry alternative alleles at the epistatic loci using a Double Generalized Linear Model (DGLM)^37^, as suggested in Rönnegård *et al.*^14^. This allowed us to simultaneously model the effects of every locus on the phenotypic mean and variance. We fitted a DGLM with linear predictors for both ean and variance as 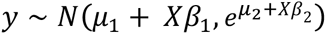 using the R-package *dglm*^38^, where *y* is the phenotype, *X* is the genotype, *β*_1_ is the effect on the mean, and *β*_2_ is the effect on the variance. Coding the genotypes in *X* as 0 and 1, β_1_ then describes the difference in mean whereas 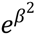 describes the fold difference in variance between the segregants with alternative alleles at the locus.

### Estimating the capacitating effects of the hub-QTL in the epistatic networks

QTL interacting with 5 or more other loci were defined as hubs. We estimated the capacitating effects of all hub-QTL as follows. For each network containing a hub-QTL, we divided the segregants into two groups based on their genotype at the hub. We then fitted the mixed model *y* = *Xμ* + *a* + *e* separately for each group. Here, *y* is a column vector containing the phenotypes, *X* is a column vector of ones, *μ* is the overall mean, 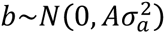 and 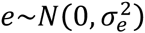. *A* is the additive kinship matrix, giving the fraction of the genome shared between each pair of segregants 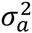 is the additive genetic variance captured by the markers, and 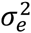 is the residual variance. *A* was calculated using the *ibs* function in the R-package GenABEL. We used the GenABEL function *polygenic* to fit the mixed model. The narrow sense heritability in each group was calculated as the intra class correlation 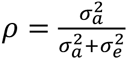.

We performed a permutation test to obtain the significance of the difference *ρ*_1_ – *ρ*_2_ between the two groups of segregants in each network. For each of the 20 traits, we randomly divided the population into two groups and estimated *ρ*_1_ and *ρ*_2_ in these as described above. This was repeated 1000 times per trait to obtain 20 empirical NULL distributions. The difference *ρ*_1_ – *ρ*_2_ was considered significant at a multiple testing threshold of 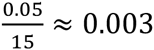.

### Quantifying non-linear effects after capacitation

To quantify the potential multiplicative action of capacitated radial alleles, we compared the fit of an additive (*y* = *μ* + *xβ* + *e*) and an exponential 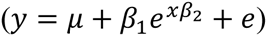 model. Here, *y* is the phenotype, *x* is the number of growth decreasing radial alleles, and *e* is the residual variance.

### Modeling the phenotypes for individual multi-locus genotype classes

For each six-locus epistatic network, we estimated the phenotypes for the 64 individual genotype classes in each network using three different models including i) additive effects, ii) additive effects and pairwise interactions and iii) additive, pairwise and three-way interactions, respectively. For ii) and iii), the included interaction-terms that, together with all dditive terms, minimized the *AIC* = 2*k* – 2*ln*(*L*). Here, *k* is the number of included parameters and *L* is the maximum likelihood value for the model. We used the R-function *step*in the *stats* package to perform the backward elimination^34^. The performances of the final models (bias and accuracy) were evaluated using 10-fold cross-validation, where the variable selection for i) and ii) above was performed within the training data in every fold.

Within each of the 64 genotype-classes, defined by the six loci in each network, we calculated the CV estimation errors as 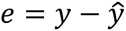. Here, *y* is the actual and 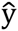 the estimated phenotype. We tested if *e* significantly deviated from 0 using a t-test. If the deviation was significant at a multiple testing threshold of 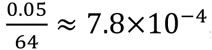, we considered the estimate for that particular genotype-class biased. The accuracy of the estimates was measured by *e*^*2*^, and for each genotype-class, we tested the difference in *e*^*2*^ between models with and without interaction-terms using a t-test. If the *e*^*2*^ was significantly lower for the interaction model, at a multiple testing threshold of 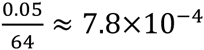, we considered these estimates more accurate.

## Simulations

In the simulations, we used the phenotypic means *μ*_1_ … *μ*_64_ in each of the 2^6^ = 64 classes for each of the 15 six-locus networks, and the total phenotypic variance 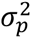 for each trait, obtained in the analyses above as a representation of the genetic architectures of these traits. In every simulation, we generated populations with the same number of segregants as in the original dataset (*n* = 4,390). The number of segregants in each genotype class was determined by the allele frequencies *p*_1_ … *p*_6_ at the six loci. For example, the number of observations with genotype *ABcDef*, where the big/small letters indicate the alternative alleles at the six loci, would be *p*_1_×*p*_2_×(1 – *p*_3_)×*p*_4_×(1 – *p*_5_)×(1 – *p*_6_×*n*. To evaluate the effect of different combinations of allele-frequencies at the loci on the results, we simulated populations with *p*_*k*_ ∈ {0.05,0.20,0.35,0.50,0.65,0.80,0.95}, where *k* = 1 ⋯6. This leads to, in total, 7^6^ allele frequency combinations. The phenotypes for the individuals in each genotype class were then simulated as *y*_*k*_~*N*(*μ*_*k*_,*σ*^2^), where *k* = 1 ⋯64.

As a comparison, we also simulated populations where the genetic architectures (i.e. the phenotype for each of the 64 multi-locus genotypes) for the 15 networks were given by the estimates obtained from a six-locus additive model fitted to the respective loci. The linear model used was *y* = *Xβ* + *e*, where *y* is a column vector containing the phenotypes, *X* is a 4,390×7 matrix, the first column consisting of ones and columns 2-7 of the genotypes of the six loci, *β* is a 1×7 column vector with the intercept and the additive genetic effects, and *e* is the residual variance. Using this model, estimates 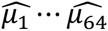 were obtained for each genotype class. The simulations were then performed across the different combinations of allele-frequencies as described above, with phenotypes given by 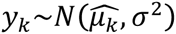.

The additive genetic variance contributed by each locus was estimated as the R^2^ from the linear model *y* = *Xβ* + *e*, fitted to the respective subset of the 7^6^ simulated populations where the allele frequency of the locus itself was 0.5 (*p*_1_ = 0.5), and the frequency at the other five loci varied in *p*_*k*_ ∈ {0.05,0.20,0.35,0.50,0.65,0.80,0.95}, where *k* = 2 ⋯6. In this model, *X* is a column vector with the genotype of the locus whose allele frequency is 0.5. The additive genetic variances were estimated in such subsets to analyze the effect of the individual locus across variable genetic backgrounds for the other loci, without the estimates being influenced by changes to the allele frequency of the analyzed locus itself.

## Author contributions

Analyses were designed by S.K.G.F., J.S.B., M.J.S., L.K. and Ö.C. Analyses were conducted by S.K.G.F and Ö.C. The manuscript was written by S.K.G.F. and Ö.C. and incorporates comments by J.S.B., M.J.S. and L.K.

## Materials and Correspondence

Correspondence should be addressed to Ö.C (email). The genotype and phenotype data are available as Supplementary information in the original publication of this data^3^.

## Supplementary text

### The additive effects and variance-heterogeneity of the hub-QTL depends on the genotypes at the radial loci

The average phenotypes for segregants with the BY allele in the IAA-network range from being very similar to that of the segregants with the RM allele at the hub-QTL, when combined with growth-increasing alleles at all five radial QTL, to being considerably lower when combined with several growth-decreasing alleles at the five radial loci (Figure 2B main manuscript). This illustrates several properties of the connection between the functional contribution by the hub-QTL to growth, and how this contribution will be represented in a statistical analysis based on quantitative genetics models.

In a classic quantitative genetics model, the main effect of the hub-QTL is modeled as an additive allele-substitution effect. The additive effect is estimated as the difference in average phenotype between the groups of segregants in the population that carry the BY allele (green) and RM allele (grey) at this locus. In a population where the BY and RM alleles of the hub-QTL segregate, but the radial loci are fixed for IAA-resistant alleles, there will be no, or a very small, difference in the average phenotype between the groups of segregants carrying the RM and BY alleles (the two leftmost boxplots in Figure 2B). The hub-QTL will thus in this population have an additive effect close to zero. By contrast, in a population where the BY and RM alleles segregate, but the radial loci are fixed for IAA-sensitising alleles, there will be a very large difference in average phenotype for the segregants carrying the RM and BY alleles (the two rightmost boxplots in Figure 2B). This will result in a very large additive effect of the hub-QTL, resulting from the total sensitising effects of all six loci in the network. The additive effect of this hub-QTL does thus not represent merely a property of this individual locus, but rather to which extent it has capacitated the effects of segregating alleles at the radial QTL in the network.

In a quantitative genetics model, the variance-heterogeneity effect of the hub-QTL is modeled as the difference in phenotypic variance between the groups of segregants that carry the BY and RM alleles at the hub-QTL. In a population where all six loci segregate, there is a large genetically controlled phenotypic variation among the segregants carrying the BY allele at the hub QTL. This is from the BY allele capacitating the genetic effects of the radial-QTL, as discussed above. The phenotypes of the segregants that carry the RM allele are, however, all very similar as the RM allele canalizes the effects of the radial loci. In such a population, there will thus be a large genetic variance-heterogeneity between the BY and RM alleles. In a population fixed for any combination of alleles at the radial loci with less than three growth-decreasing alleles, there is almost no difference in phenotypic variance between the groups of segregants that carry the RM and BY alleles at the hub-QTL. Hence, also the marginal variance-heterogeneity effect at the hub-QTL in a quantitative genetic analysis does not only represent a property of the individual locus, but is rather highly dependent on the ability of the hub-QTL to capacitate the effects of the radial QTL in the network.

Our results thus empirically illustrate how the estimates obtained for the main effects (here additivity and variance-heterogeneity) from a quantitative genetics model do not represent how the locus independently contributes to trait variation. Rather, they quantify the total contribution of the locus in the genetic context of the analyzed population.

### Some radial networks contain hub-QTL with known polymorphic candidate genes

For two networks, regulating growth on media containing high levels of Copper and Manganese, the respective hub-QTL map to genes that are polymorphic between BY and RM and involved in heavy-metal ion detoxification. The hub-QTL for growth on medium with Copper Sulfate maps to *CUP1,* a copper-activated metallothionein whose copy number is known to vary between the BY and RM strains^1^. The BY strain possesses fewer copies of *CUP1*and as reported previously^2^, this leads to greater copper ion tolerance. Here, however, only segregants with three or more sensitizing alleles at the radial loci, and the *CUP1*_*RM*_ allele, grow more poorly than segregants with the more robust *CUP1*_*BY*_ allele (Figure 3D). A hub-QTL for growth on medium with Manganese sulfate maps to the *PMR1* gene, encoding an active Mn-ion transporter, for which the *PMR1*_*RM*_ allele confers manganese resistence^3^. The multi-locus genotype-values in the network were consistent with this as segregants with a *PMR1*_*BY*_ allele grow poorly regardless of which alleles they have at the radial loci (Figure 3B). However, alleles at the radial loci can also contribute to a sensitization of the yeast to Manganese, illustrated by the observation that segregants with the *PMR1*_*RM*_ allele and low growth alleles at all five radial loci grow almost as poorly as those with the *PMR1*_*BY*_ allele. (Figure 3B). No obvious functional connections between the genes in the radial-QTL and the hub were found, for any of these networks. One hub-QTL affecting growth on medium with Cobalt chloride maps to *HAP1*. This gene is a known key regulator of transcription in crosses between the BY and RM strains. Cobalt stress is often used to mimic hypoxia^4^, and *HAP1* encodes a transcription factor involved in the transcriptional responses at many genes to levels of heme and oxygen^5^. The *HAP1*_*BY*_ allele has been disrupted by a transposon, causing a loss-of-function for some *HAP1* functions, but not all^5^. Here, segregants with the *HAP1*_*BY*_ allele were less responsive to the positive effects on growth of the alleles at the interacting radial loci, which is consistent with a stronger transcriptional activation of other genes in response to Cobalt stress in segregants with the non-disrupted *HAP1*_*RM*_ allele. Our finding of a radial epistatic network around a key transcriptional regulator is coherent with earlier work showing that molecular, including expression, networks are generally heavy-tailed^6^, i.e. contain few hubs and many low-degree nodes (Figure 1A).

---

**Supplementary Figure 1.**
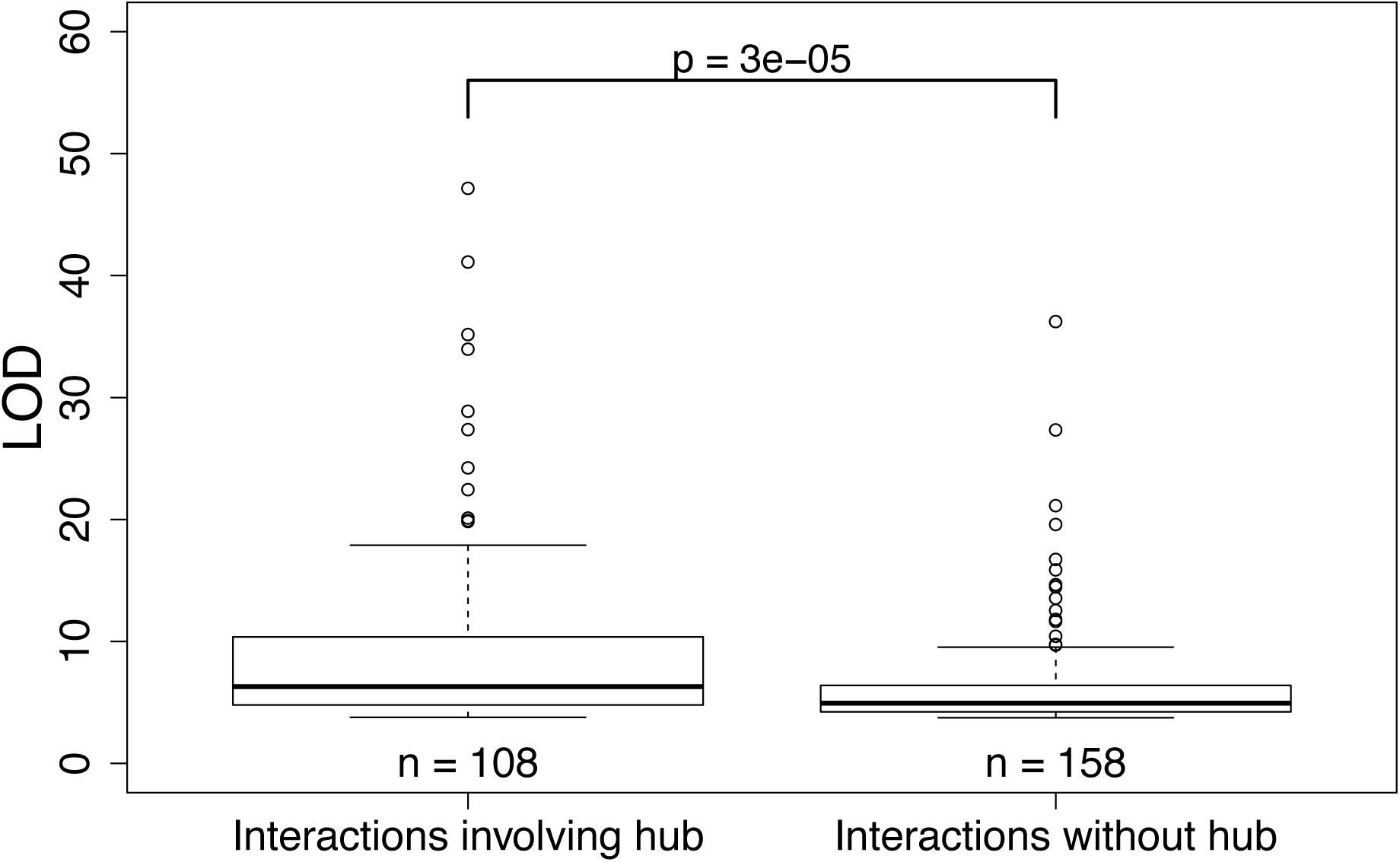
LOD-scores for all significant genetic interactions (n = 266) involving the 330 epistatic QTL detected in this population. The left box-plot displays the LOD-scored for the genetic interactions between hubs (involved in more than 4 interactions) and radial loci. The right box-plot displays the LOD-scores for the genetic interactions the do not involve a hub.

**Supplementary Figure 2.**
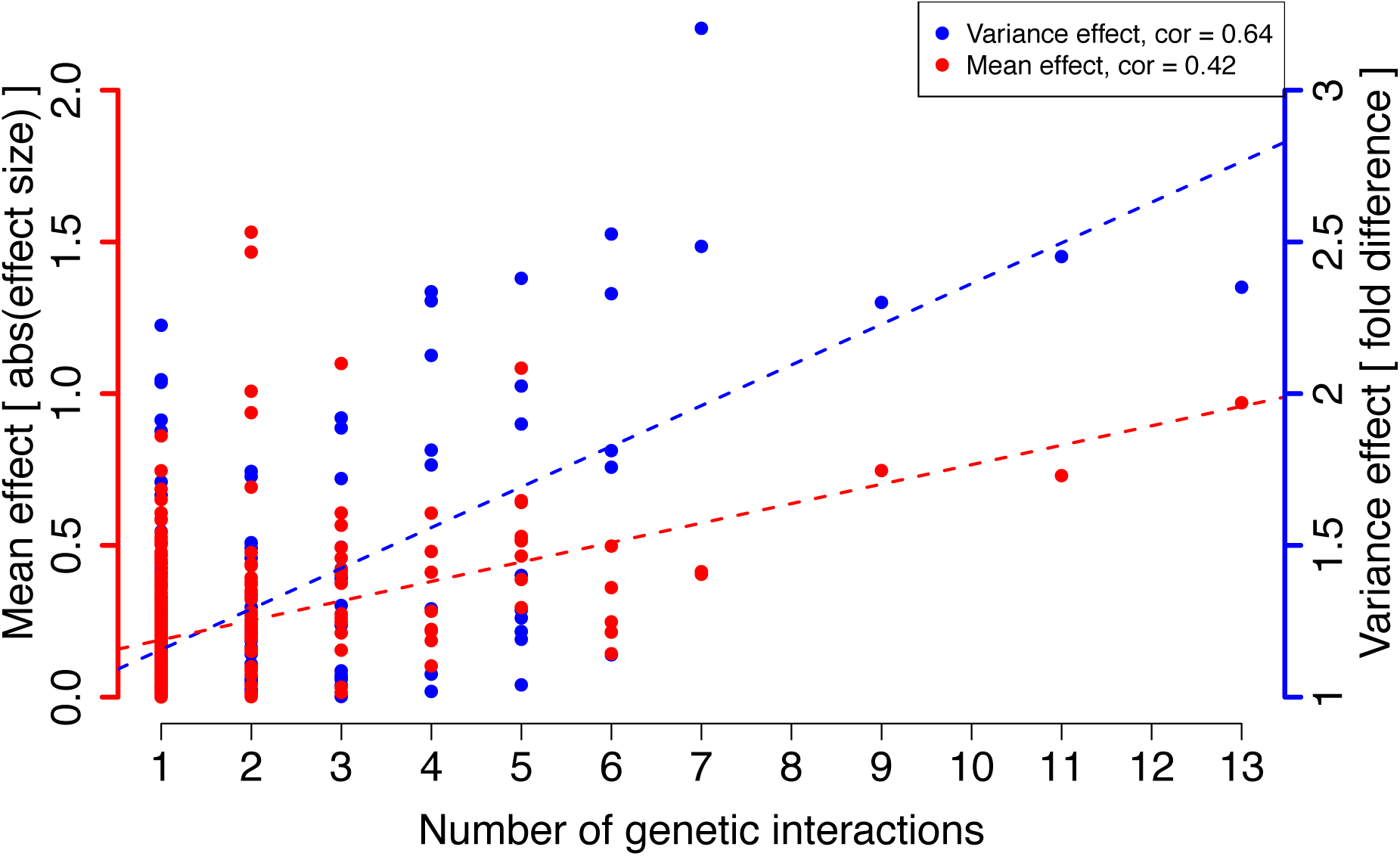
The correlations between the number of genetic interactions that the loci are involved in and their marginal additive and variance-heterogeneity effects. The X-axis shows the number of pairwise genetic interactions each locus is involved in. The left Y-axis shows the marginal additive and the right Y-axis the marginal variance-heterogeneity effect on the phenotype. Each dot in the figure represents a locus. The red dots show the marginal effect on the mean of the phenotype (cor = 0.42; p < 1 x 10^−12^), whereas the blue dots show the effect on the phenotypic variability (cor = 0.64; p < 1 x 10^−12^).

**Supplementary Figure 3.**
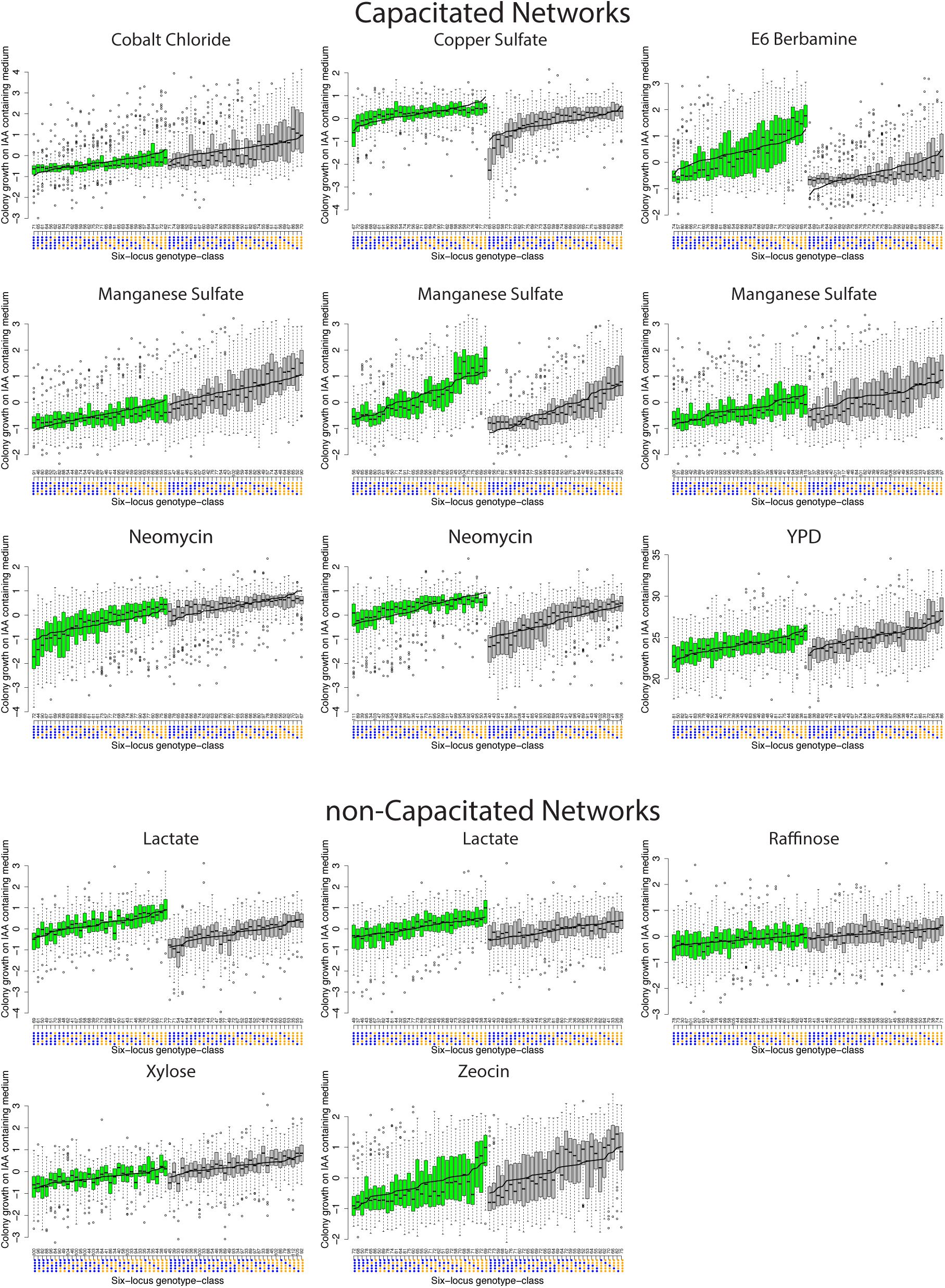
Phenotypic distributions in segregants with different combinations of alleles across the six loci in 14 epistatic networks affecting growth in different media. The networks are divided depending on whether the hub-QTL is a significant capacitor or not. One Tukey-boxplot is provided for each of the 64 genotypic classes in every network. The color gives the genotype at the hub-QTL (Green/grey boxes for BY/ RM alleles). The x-axis represents the six-locus genotype class, where blue/orange dots indicate growth-increasing/decreasing alleles at the five radial-QTL in the network. The black lines through the boxes illustrates the additive model-based estimates of the phenotypes for the 64 genotype-classes. The number above the x-axis shows the number of segregants in each genotype class.

**Supplementary Figure 4.**
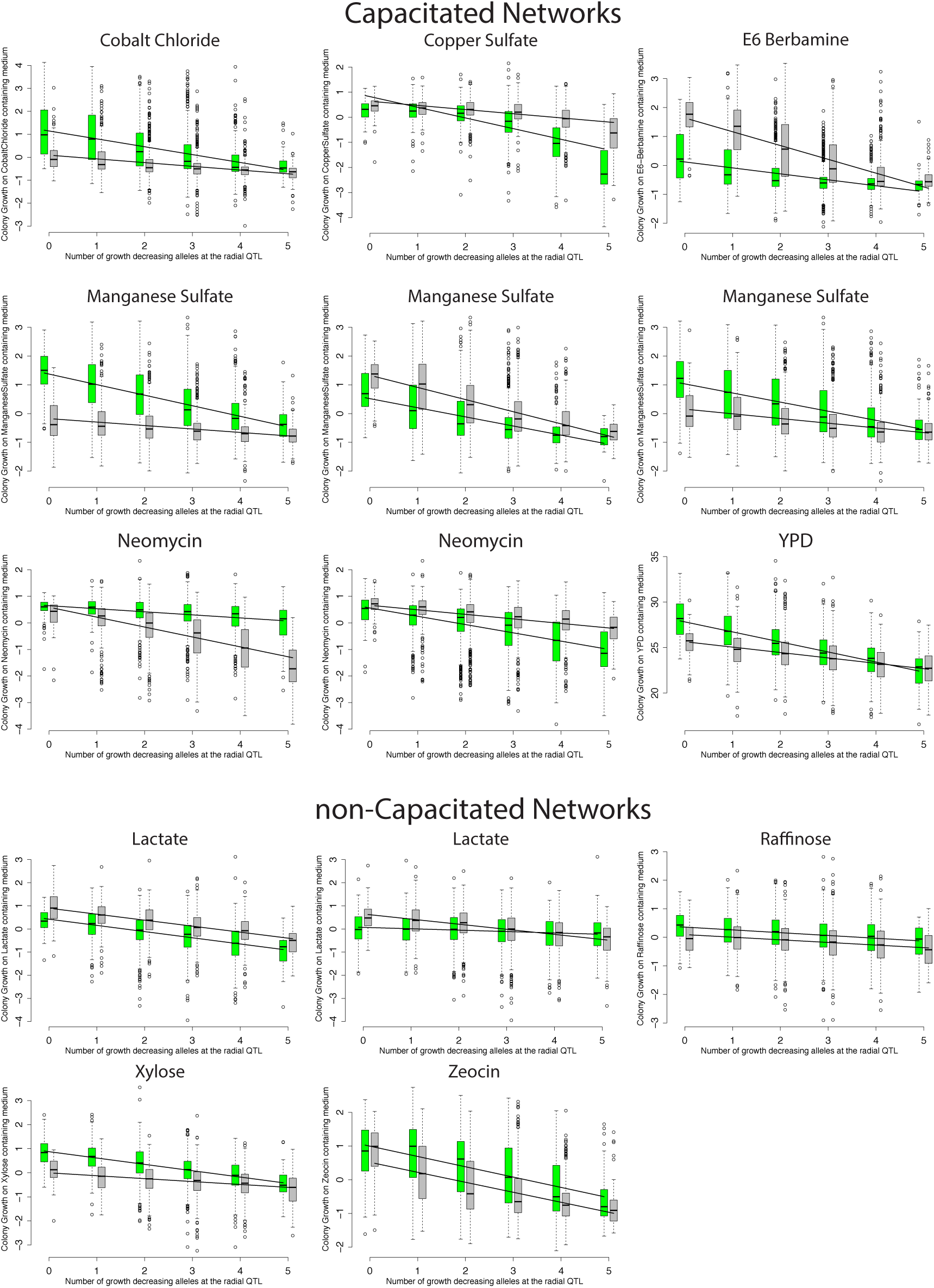
Phenotypic distributions in segregants with varying number of growth decreasing alleles at the radial loci across the six loci in 14 epistatic networks affecting growth in different media. The networks are divided depending on whether the hub-QTL is a significant capacitor or not. Each Tukey boxplot represents a group of segregants that share the same number of growth decreasing alleles at the five radial QTL in the respective networks. The segregants are divided and colored based on the genotype at the hub-QTL (Green/grey boxes for BY/ RM alleles). The x-axis gives the number of growth decreasing alleles at the radial QTL and the number of segregants in each group. The regression lines illustrate the fit for linear additive and non-linear exponential models, respectively.

**Supplementary Figure 5.**
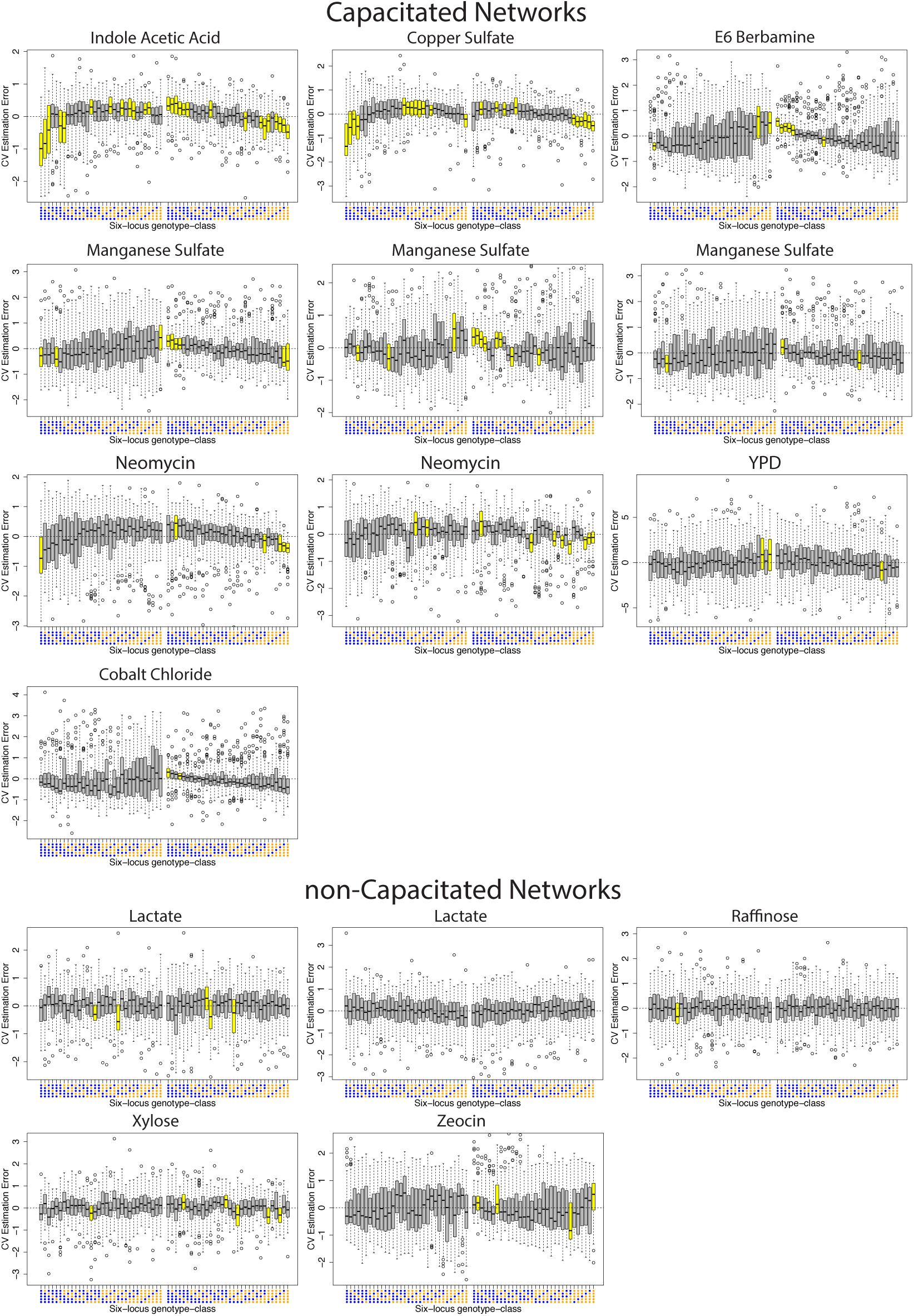
Phenotypic estimation bias for the individual multi-locus genotype-classes in the 15 epistatic networks. The networks are divided depending on whether the hub-QTL is a significant capacitor or not. Each Y-axis gives the cross-validated estimation error (bias) for the six-locus additive model representation of the genotype-value of each individual multi-locus genotype-class as compared to the actual genotype value estimated directly from the data. Each boxplot shows the distribution of prediction errors in one of the 64 genotype-classes in the network. The 32 leftmost boxplots represent the genotype-classes with the capacitor hub-QTL allele (or that with the highest h^2^ in the case of a non-significant capacitor hub). Significant biases, i.e. where the estimation errors deviate significantly from zero, are colored in yellow.

**Supplementary Figure 6.**
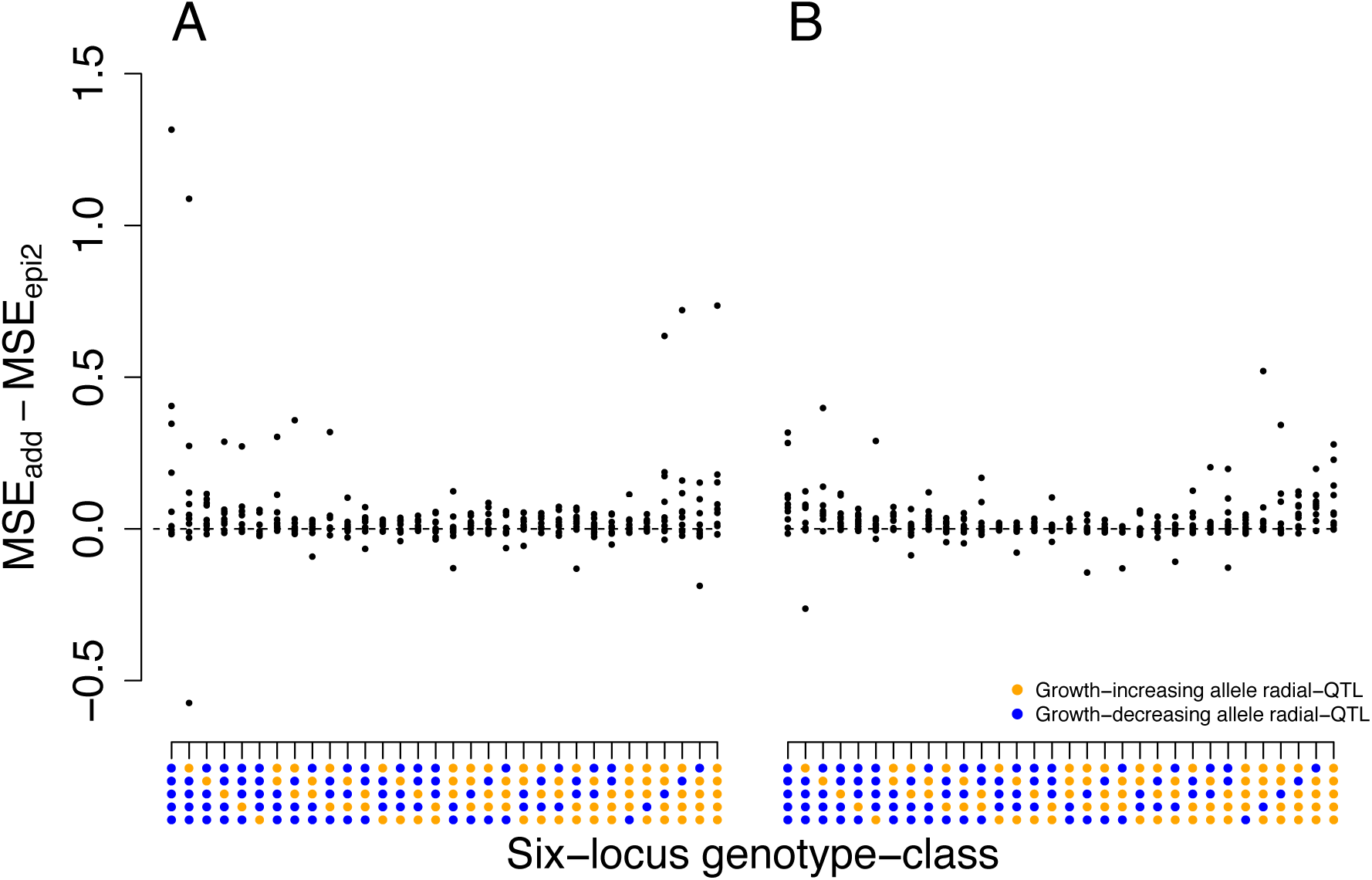
Difference in estimation accuracy between additive and epistatic models. The figure displays cross-validation results from the 10 epistatic networks where the hub-QTL is a significant capacitor. The Y-axis gives the difference in cross-validated mean squared error (MSE) between an additive model (MSE_add_), and a model including pairwise interactions (MSE_epi2_). Each dot corresponds to the difference in MSE for one genotype class. The difference is measured as MSE_add_ − MSE_epi2_, meaning that positive values correspond to genotype classes where the estimation accuracy is improved when using an epistatic model.

